# Nanoscale investigation in 3D scaffolds of cell-material interactions for tissue-engineering

**DOI:** 10.1101/383117

**Authors:** Donata Iandolo, Fabrizio A. Pennacchio, Valentina Mollo, Domenico Rossi, David Dannhauser, Bianxiao Cui, Roisin M. Owens, Francesca Santoro

## Abstract

Cell fate is largely determined by interactions that occur at the interface between cells and their surrounding microenvironment. For this reason, especially in the field of cell- and tissue-engineering, there is a growing interest in developing characterization techniques that allow a deep evaluation of cell-material interaction at the nanoscale, particularly focusing on cell adhesion processes. While for 2D culturing systems a consolidated series of tools already satisfy this need, in 3D environments, more closely recapitulating complex *in vivo* structures, there is still a lack of procedure furthering the comprehension of cell-material interactions. Here, we report for the first time the use of a SEM/FIB system for the characterization of cellular adhesion in 3D scaffolds fabricated by means of different techniques. Our results clearly show the capability of the developed approach to finely resolve both scaffold-cells interface and nanometer scale features of cell bodies involved in the upregulation of cellular behavior. These results are relevant for studying cellular guidance strategies and for the consequent design of more efficient cell-instructive platforms for tissue-engineering applications as well as for *in vitro* 3D models.

## Introduction

The adhesion between cells and biomaterials plays a key role in the success of many biomedical platforms such as prosthetics, artificial organs and auxiliary devices^1^. In particular for implantable devices, optimal adhesion is crucial to avoid the implant rejection and then to ensure its complete integration into the body^2^. Such processes are regulated by the interaction of single cells with the biomaterial itself^3,4^. For this reason, in the last decades, many efforts have been carried out to properly engineer the material surface and bulk properties, *i.e.* surface topography^5^, chemical functionalization^6^, material stiffness^7,8^ for delivering complex sets of cell-instructive signals capable to specifically affect cell fate^9,10^. The final goal in designing such platforms is in fact to create instructive adhesion areas capable of guiding in a controlled way several cell functions fundamental for the good implant integration, such as migration, proliferation, differentiation and the synthesis of endogenous extracellular matrix (ECM). Here, particular attention has been dedicated to the development of 3D systems that better replicate the structural organization of living tissues and that, compared to 2D platforms, have been extensively demonstrated to affect cellular behavior in a more realistic way^11,12^. For instance, 3D matrices/scaffolds have been fabricated with different degrees of structural organization to correlate *in vitro* cells responses with different and well-defined cell-material interactions^13–16^. In this context, in order to evaluate the effect of specific functional features upon cellular response, one can get fundamental informative cues from the investigation of the interface between cells and 3D biomaterials at the macro, micro and nanoscales.

Optical microscopy is effective in providing accurate characterization of cellular features, *i.e.* nucleus, actin filaments, focal adhesion, with a resolution down to the sub-micrometric scale (hundreds of nm) through direct imaging by using fluorophores^17,18^. However, electron microscopy is the ultimate technique to achieve highest resolution (down to a few nanometers) in the investigation of the cell-material interface because of its capability to resolve features with the highest resolution (down to a few nanometers). In combination with the fluorescence-based techniques, it could in fact substantially widen the overview on cell-material interaction adding important information related to cellular architecture in response to specific cell-instructive signals^19^. While traditional electron microscopy specimen preparation well fits 2D cell-material systems^20^, it finds major limitations in the case of 3D matrices^21^. In resin embedding-based procedures coupled with mechanical sectioning, removal of the support material is required and it is achieved *via* physical etching. This procedure can induce artefacts particularly at the contact area between cells and the material. While this is a common approach for cells on 2D materials, it is obviously incompatible with 3D materials since cells can grow in all directions and the removal of the material itself could cause collapse of the cellular components. In light of this, the ideal approach is the one allowing maintaining and directly sectioning the support materials in place with the cells. Here, resin-embedded specimens combined with support material can be alternatively sectioned *via* focused ion beam (FIB)^22^. However, also in this case, the presence of a large resin matrix does not allow the selection of a region of interest (ROI) while the whole specimen has to be sectioned^23,24^. Other methods have been established to characterize the interaction of 3D scaffold-like materials with cells involving hard drying procedures of specimens combined with scanning electron microscopy (SEM)^25,26^. However, typical artifacts such as cracks and cell detachment can be visible and this is not representative of the effective cell-material interface. Effectively, there is a lack of procedures which allow preserving both the 3D material structure and the cells in place and to, subsequently, selectively perform sectioning for high-resolution investigation of the cell-3D material interface.

Recently, ultra-thin plasticization (UTP) of cells has been successfully performed to overcome the aforementioned limitations in case of adherent cells on 2D and pseudo-3D materials (surface with protruding micro and nanostructures)^27–29^. This technique allows the allocation of a region of interest (ROI) for a selective cross sectioning *via* FIB and a high resolution imaging (5-10 nm) *via* SEM, such that plasma membrane as well as intracellular compartments can be visualized^30^.

Here, we present the implementation of an UTP procedure for ultimately studying the interface between 3D polymeric architectures and cells at the nanoscale, by means of electron microscopy technologies. Moreover, we show our capability in coupling such analysis with confocal characterization for the evaluation of cell-material 3D interactions (*i.e.* adhesion, proliferation, differentiation). In particular, we present two 3D scaffold types which differ by spatial arrangement of their backbone (ordered *vs.* non-ordered) and their interaction with cells, showing the capability of the implemented technique in resolving structural features with nanometer resolution. In fact, the acquired SEM images allow for whole-cell imaging, while FIB-mediated sectioning reveals, in detail, how the plasma membrane and intracellular organelles differently distribute in response to the scaffold geometry.

First, we fabricated polymeric scaffolds with two different geometries. By means of a 2-photon polymerization lithography system (Nanoscribe Photonic Professional GT, Nanoscribe GmbH), ordered scaffolds have been obtained. A 4-layer example is shown in the scanning electron micrograph in **Figure 1A**. Structures were fabricated processing a commercial negative photoresist using a constant power of 60 mW and a scan speed of 1000 μm/s. The design consists of adjacent “cages” with a ~50 μm openings/height. 8×8 cages have been first serially stitched in x,y direction to form the individual layer (**Figure 1B**), that was in turn connected in the z-direction to form the entire structure. Adjacent cages were stitched together considering an overlap of ~1 μm to ensure the necessary final structural stability. This serial fabrication approach allows us to obtain very complex and stable structures that would be harder to fabricate using a one-step fabrication process. Thanks to this strategy, it is also possible to change the overall dimensions of the structure without the need to recur to complex structural design iterations. The scaffolds were then sterilized and coated with a fibronectin solution to encourage cell adhesion. Human glioblastoma astrocytoma (U87) cells were finally seeded on the ordered scaffolds and prepared for the successive analysis (confocal and SEM/FIB analysis).

**Figure 1:**
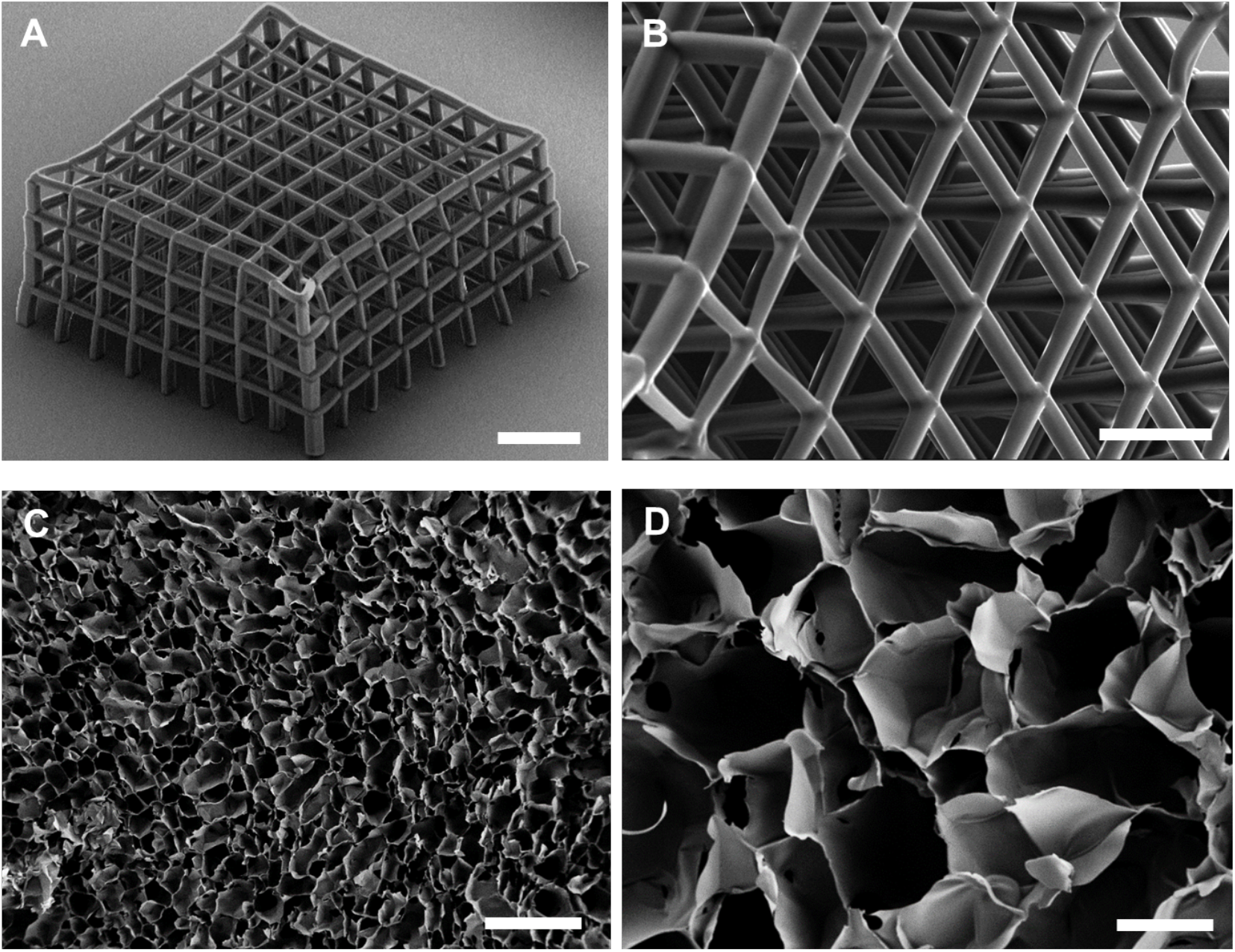
SEM characterization of fabricated scaffolds. A-B) Ormocomp ordered scaffolds fabricated by means of 2-photon patterning. C-D) PEDOT:PSS scaffolds prepared by the ice-templating technique. Scale bars: A) 150 um, B) 50 um, C) 400 um, D) 100 um, inset D) 200 um.

Non-ordered PEDOT:PSS scaffolds were fabricated, instead, *via* the so-called ice-templating technique^25,31^. Macroporous scaffolds were prepared following a slightly modified version of the procedure reported in previous studies^25,31^. An aqueous dispersion of PEDOT:PSS was prepared as previously reported^25^ (described in the **Experimental Procedure**), frozen at a specific rate and then the ice crystals were let to sublime under controlled conditions giving rise to a highly porous structure. 400 μm thick slices were prepared using a Vibrating Blade Microtome (LEICA VT1200) and subsequently used for seeding human Adipose Derived Stem Cells (hADSC, Lonza). The ice-templating technique creates structures with interconnected pores displaying a broad range of diameters as shown in **Figure 1C&D**. The obtained cavities are irregular and, as it is possible to appreciate from the exemplary image in **Figure 1D**, pores have diameters in the range 100-250 μm facilitating cells infiltration and media penetration. Cells were seeded taking care that the cell suspension was distributed homogeneously throughout the sample surface.

After cell culture, specimens were first observed *via* confocal microscopy (**Figure 2B&E**) to assert cells adhesion and proliferation onto the investigated surfaces and then by SEM-FIB microscopy coupled with the developed UTP procedure (**Figure 2A**). As shown in **Figures 2C&D and figures 2F&G**, no resin excess is present on the specimens such that both the material surface and individual cells are distinguishable. This applies to UTP preparations performed with the two investigated embedding resins: EPON and SPURR resins. As a reference, viscosity measurements were run for both EPON and SPURR resins (**Supplementary Information S1**) after 0, 30, 105 days from preparation. Such measurements were performed at room temperature with a rotational rheometer. Importantly, as reported from the viscosity measurements, these resins experience an increase in their viscosity with time, hindering the removal of the excess material in the final steps of the infiltration. It was thus crucial to proceed with samples preparation starting with freshly prepared resin mixtures.

**Figure 2:**
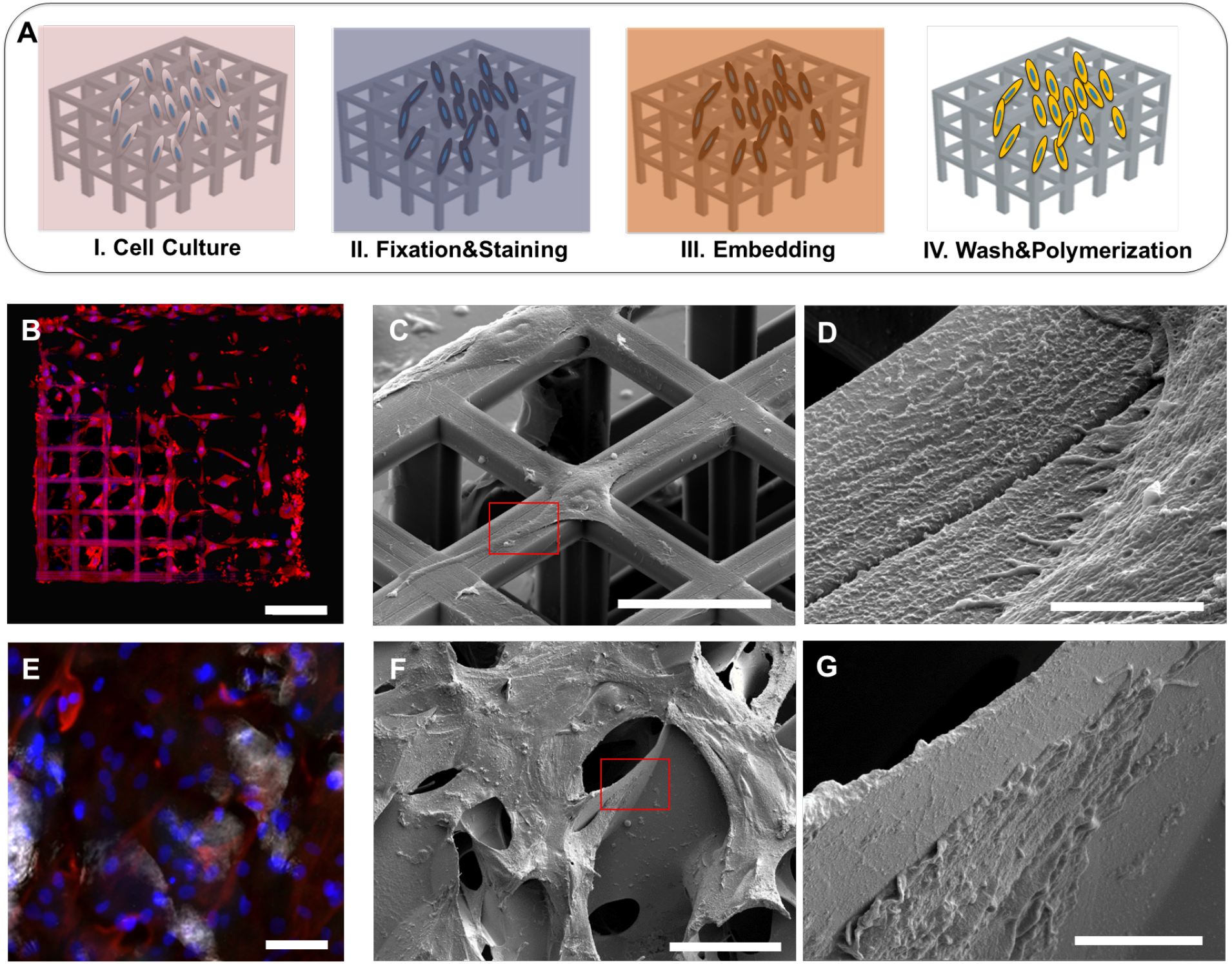
SEM characterization of plasticized cells on scaffolds. A) Schematic of UTP process on scaffolds. B) U87 glioma cells spreading on ordered scaffold with fluorescence staining of actin (magenta) and nucei (blue). C-D) SEM images of plastified U87 on ordered scaffold. E) hADCs spreading on non-ordered scaffolds with fluorescence staining of vimentin (purple) and nuclei (cyano). F-G) Plastified hADCs on non-ordered scaffolds. Scale bars: B) 200 um, C) 40 um, D) 3 um, E) 50 um F) 100 um, G) 10 um.

Then, we first characterized cell-scaffold interaction with confocal imaging, where features like cytoskeleton architecture and nuclei distribution could be assessed (**Figure 2B&D**). Successively, we performed the UTP procedure for electron microscopy characterization. Here, we observed cell-material 3D interaction both at the macroscale visualizing the cell layer conformation on a large ROI (**Figure 2C&F**) and, then, at higher magnification (micro- and nanoscale) selectively imaging cell-material point of contacts (**Figure 2D&G**). From these images, we were able to easily evaluate how cells colonize and interact with the structural features of the ordered scaffolds. In particular it is possible to clearly and directly observe cell alignment and stretching on the scaffold surfaces and how they wrap their plasma membranes around the 3D structures. Cells seeded on the macroporous scaffolds penetrate into the pores and deposit an abundant ECM layer during their proliferation. Interestingly, its composition in terms of protein and glycosaminoglycans (GAG) content as well as types of present proteins and its importance for cell differentiation has just been investigated in 2D culture conditions ^32^. The presence of a very thin layer of resin, as shown in the images, preserves cell integrity at different densities, then leading to high quality observation of both sparse cells distributed on ordered scaffolds as well as denser, more complex cells architectures such as those on the non-ordered scaffolds. Furthermore, for both scaffold types it is possible to resolve ultrastructures and cellular protrusions such as filopodia, fundamental features involved in the formation of focal adhesion complexes (**Figure 2D&G**).

As mentioned earlier, when in contact with 3D scaffolds, cells show structural conformations which differ from those typically displayed by cells adhering onto 2D surfaces^33^. These different types of interactions, moreover, have been demonstrated to deeply influence cells tensional state and, as a consequence, their behavior^12,33,34^,. Depending on the scaffolds architecture, cells can partially adhere to the local surface area or even be suspended across the pore bridging two sites of the support material. These conditions reflect high tensional states of the membrane which could often result in cracks when specimens are prepared with hard drying techniques^16^. This aspect limits the possibility to fully characterize cell-material interactions in complex cellular 3D systems and then the capability to properly evaluate the effect of a peculiar design on cell adhesion processes.

The developed procedure allowed us to access and characterize cells positioned in locations inaccessible with other approaches^16^. Here, considering a top view of an ordered scaffold with cells, we were indeed able to analyze 3 different relative positions of adherent cells in respect to the arms of the ordered scaffold: cells adherent to the strut surface (**Figure 3A**), cells suspended between two struts (**Figure 3C**), cells partially attaching to one arm (**Figure 3E**). To further characterize the effect of structural features upon cell adhesion, we performed a FIB-based sectioning of both material and the cell body. Once ROIs were located, these were coated with a platinum layer first by electron beam-assisted deposition and then by ion beam-assisted deposition^28^ to reach a final thickness of ~1μm (**Supplementary Information S2**). Subsequently, the material close to the ROI was removed by trenching out scaffold areas with a depth of 10 μm (nominal, as for silicon) as shown in **Supplementary Information S3**. After the second round of metallization is performed in the section of interest (**Supplementary Information S4**) that had been previously created perpendicularly to the main direction of the square arm, a final polishing was performed by using a low ion beam current (~ 0.7 nA-80 pA). The resulting polished sections are showed in **Figure 3B, D, F**. Remarkably, cellular ultrastructures, nuclei and plasma membrane have been resolved. (**Figure 3G-H**). Moreover, 5-50 nm vesicular invaginating processes are visualized, resembling inwards buddings as previously reported ^28,30,34^ (**Supplementary Information S5**).

**Figure 3:**
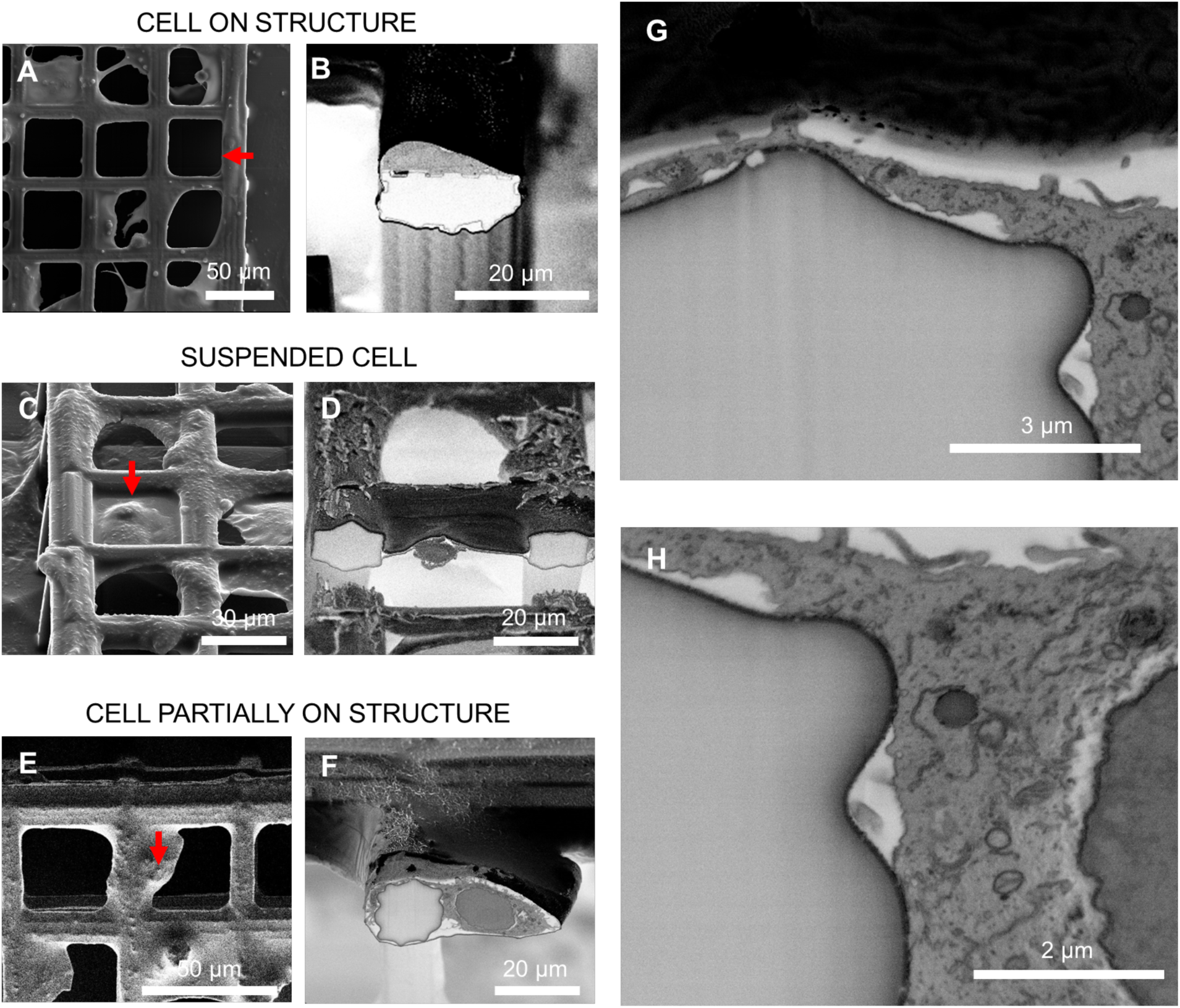
Cross sectioning of cells on ordered scaffolds. A) Top view of a cell growing on top of a square arm; B) 52° tilt view of cross section from position selected in A; C) 52° tilt view of cell suspended and spreading between two arms; D) 52° tilt view of cross section from position selected in C; E) Top view of a cell partially attaching one arm and spreading over the perpendicular direction; F) 52° tilt view of cross section from position selected in E; G-H) Zoom-in of cross areas where cell located in E-F. A, C, E, G are acquired in secondary electrons mode. B, D, F are acquired in backscattered mode and inverted.

Finally, the sectioning procedure has been performed on specimens with cells grown onto the non-ordered PEDOT:PSS scaffolds. In this case, cells freely colonize the scaffold surface and penetrate in the pores, depositing an abundant layer of ECM (Figure 4), necessary for cell adhesion, survival and cell to cell communication as well as to adapt to the culture substrate properties^32,35^. Compared to the case of ordered scaffold it is worth noting that in this case, because of both the irregular scaffold architecture and the complex cellular rearrangement, the most advanced microscopy techniques find greater limitations in imaging the scaffold core ^16^. While for ordered scaffolds it is possible to create a cross section following the geometry of the 3D structure and the relative position of cells on it, in the case of non-ordered scaffolds the region of interest is located mainly based on the upmost layer of cell visible *via* SEM (**Figure 4A**). Also in this case, as shown in figure 4B, it is possible to perform nm sectioning and clearly observe the interface of multiple cells with the underlying scaffold for the evaluation of the adhesion processes. Interestingly, the developed approach allowed the resolution of details of the ECM deposited by cells, giving also the possibility to easily distinguish different ECM elements like laminin, fibronectin and collagen^36^ and its distribution (**Figure 4B-4C**). Additionally, from such images also cells spatial rearrangement and cell-cell interaction could be resolved (**Figure 4B**). In particular, the brighter areas in **Figure 4C** reveal that residual resin is present in the inner areas, which is advantageous for preserving cell-cell position and the cellular structures as well as ECM components. Furthermore, with a particular focus on the interface contact area, also in this case it is possible to completely resolve the plasma membrane approaching the surface area of the PEDOT:PSS cavity. An example is shown in **Figure 4B** where the plasma membrane lies below a scaffold cavity and ruffles around the irregularities at the material surface.

**Figure 4:**
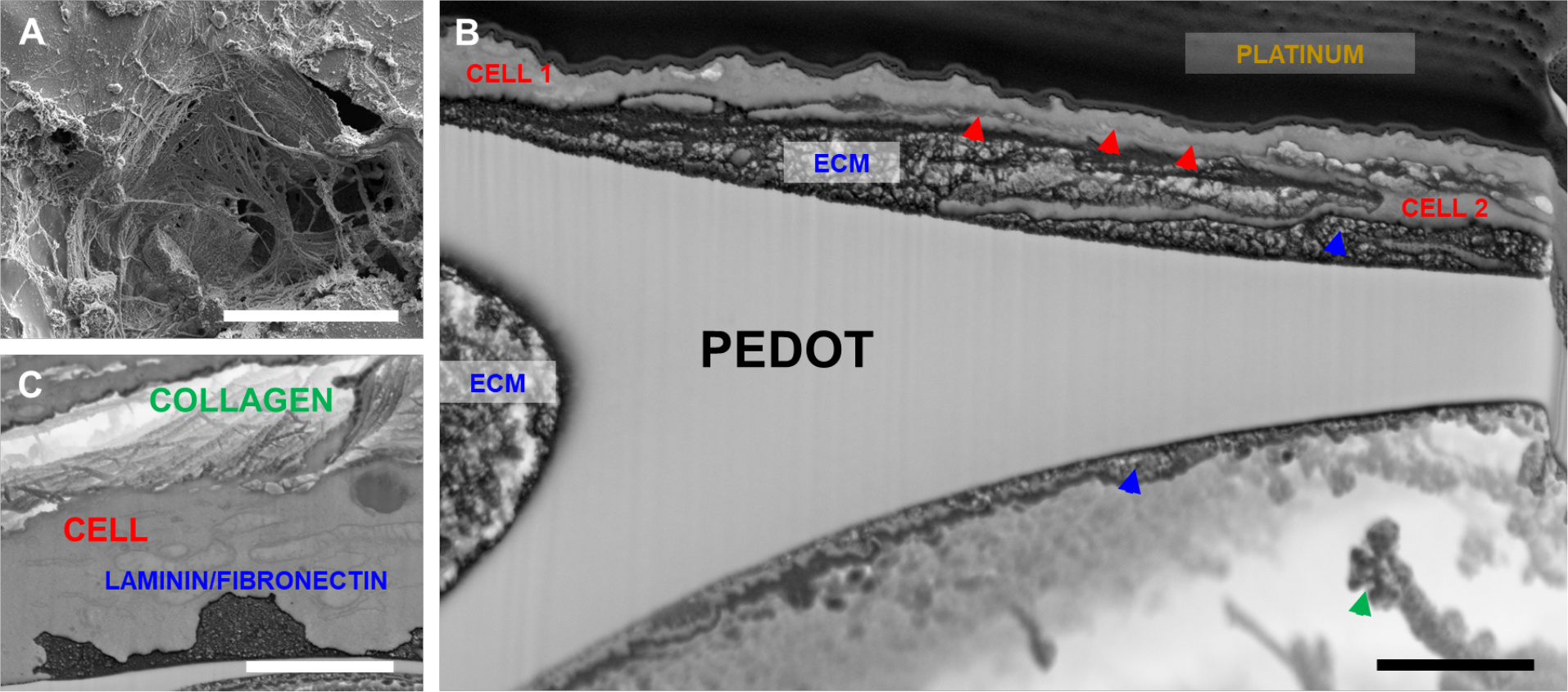
Cross sectioning of cells on non-ordered scaffolds. A) Top view of plastified adipose cells and extracellular matrix on PEDOT:PSS scaffold B-C) Cross sections reveal cellular and extracellular component. Red arrows highlight the cell-cell contact points, blue arrows highlight ECM components and green arrow highlights collagen fiber. Scale bars: A) 40 um, B) 1 um, C) 2 um.

Definitely, the unique capability to directly observe cell sections with the aforementioned resolution in 3D complex environments opens the way to a more efficient evaluation of the effects of specific instructive signals on cellular features involved in the upregulation of important biological events such as, for example, endocytosis, proliferation, migration and differentiation.

## Conclusions

We have shown two types of scaffolds for tissue engineering realized by two-photon lithography and ice-templating technique. The two fabrication methods lead to two distinct geometries, cage-like and random-distributed cavities, respectively. To investigate the influence of the different 3D scaffold architectures on cell-material interactions, we performed on the same samples both confocal imaging and UTP before imaging samples by scanning electron microscopy. Remarkably, the presented UTP procedure, by limiting the volume of resin remaining in the samples after fixation and infiltration, allows the visualization of both scaffold surface and cells with nanoscale resolution. Moreover, the heavy metal staining allowed the resolution of intracellular components such as nuclei and plasma membrane, vesicles as well as ECM components. The unique possibility to fully characterize the interface both at the micro and nanoscale allows for an accurate evaluation of the effect of the properties of different materials (surface chemistry/geometry/stiffness) on the interaction of 3D scaffolds with the investigated cells. The high versatility of the developed approach allows for the investigation of a broad range of 3D scaffolds that could differ for both structural organization and material composition, performing observations from the tissue to the single cell level. This innovative approach opens the way for a deeper comprehension of the cell-material interaction in 3D environments, that can be leveraged to rationally design more efficient new generation of tissue engineering materials and implants.

## Acknowledgments

F.S. acknowledges the Stanford Nanoshared Facility for the use of the dual beam machine Helios 600i, Michele Dipalo, and Francesco De Angelis for the use of dual beam machine Helios 650. D.I. acknowledges support from a Marie Skłodowska-Curie Individual Fellowship (IF-EF), (Grant Agreement No; 704175).

